# The genomic determinants of adaptive evolution in a fungal pathogen

**DOI:** 10.1101/176727

**Authors:** Jonathan Grandaubert, Julien Y. Dutheil, Eva H. Stukenbrock

**Author notes:** Eva H. Stukenbrock, Environmental Genomics, CAU Kiel, Am Botanischen Garten 1-11, 24118 Kiel, Germany, Julien Y. Dutheil, Research group Molecular Systems Evolution, Max Planck Institute for Evolutionary Biology, August-Thienemann-Str. 2, 24306 Plön, Germany.

## Abstract

Antagonistic host-pathogen co-evolution is a determining factor in the outcome of infection and shapes genetic diversity at the population level of both partners. While the molecular function of an increasing number of genes involved in pathogenicity is being uncovered, little is known about the molecular bases and genomic impact of hst-pathogen coevolution and rapid adaptation. Here, we apply a population genomic approach to infer genome-wide patterns of selection among thirteen isolates of the fungal pathogen *Zymoseptoria tritici*. Using whole genome alignments, we characterize intragenic polymorphism, and we apply different test statistics based on the distribution of non-synonymous and synonymous polymorphisms (pN/pS) and substitutions (dN/dS) to (1) characterise the selection regime acting on each gene, (2) estimate rates of adaptation and (3) identify targets of selection. We correlate our estimates with different genome variables to identify the main determinants of past and ongoing adaptive evolution, as well as purifying and balancing selection. We report a negative relationship between pN/pS and fine-scale recombination rate and a strong positive correlation between the rate of adaptive non-synonymous substitutions (ω_a_) and recombination rate. This result suggests a pervasive role of Hill-Robertson interference even in a species with an exceptionally high recombination rate (60 cM/Mb). Moreover, we report that the genome-wide fraction of adaptive non-synonymous substitutions (α) is ~ 44%, however in genes encoding determinants of pathogenicity we find a mean value of alpha ~ 68% demonstrating a considerably faster rate of adaptive evolution in this class of genes. We identify 787 candidate genes under balancing selection with an enrichment of genes involved in secondary metabolism and host infection, but not predicted effectors. This suggests that different classes of pathogenicity-related genes evolve according to distinct selection regimes. Overall our study shows that sexual recombination is a main driver of genome evolution in this pathogen.

## Introduction

Antagonistic host-pathogen interactions drive co-evolutionary dynamics between pathogens and their hosts. Signatures of selection in genomes inform about mechanisms of evolution and identify targets of selection at interacting loci (Möller & Stukenbrock 2017). In general, genome studies of microbial pathogens have focused on rapidly evolving genes involved in pathogenicity, such as “effector” genes encoding proteins that interfere with host defenses and may determine host range of the pathogen (Lo Presti *et al.* 2015). Effector genes of fungal pathogens have frequently been found to associate with repetitive DNA and it is proposed that repeat rich genome compartments provide particularly favorable environments for rapid evolution of new virulence specificities (e.g. (Ma *et al.* 2010; Spanu *et al.* 2010; Klosterman *et al.* 2011; Daverdin *et al.* 2012)). Repetitive DNA may locally increase mutation rate and contribute to gene duplications and structural variation among alleles. Yet little is known about the factors that shape genome evolution in fungal pathogens, in particular the interplay of mutation, natural selection, genetic drift and, for sexually reproducing species, recombination along the genome of fast evolving pathogens.

Evolution of genes involved in the antagonistic interaction with the host can be driven by positive selection whereby new alleles recurrently replace existing alleles in response to allelic changes in the host, a scenario termed arms race evolution (Van Valen 1973; Tellier *et al.* 2014). Variation in pathogenicity related genes can also be maintained by balancing selection, a trench-warfare scenario where a set of alleles are maintained in the population over long evolutionary times (Stahl *et al.* 1999). In plants, balancing selection has been described as a main driver of evolution in genes encoding resistance proteins (e.g. (Tian *et al.* 2002; Huard-Chauveau *et al.* 2013)), however the importance of balancing selection in pathogen genomes is less understood.

Population genomic data reflect signatures or past and on-going selection acting on the organism. While past signatures of selection can be related to ecological specialization, on-going positive selection reflects local adaptation in the existing population. In plant pathogens, signatures of on-going selection can reflect host-pathogen arms race or trench warfare evolution as well as adaptations to other local environmental conditions, in agricultural systems notably fungicide treatments (Hayes *et al.* 2015; Delmas *et al.* 2017). Rapid adaptation is fueled primarily by large effective populations sizes as well as high recombination and mutation rates that promote the emergence, spread and fixation of new advantageous alleles. In research of Eukaryote pathogen genome evolution, most studies have used genome scans to detect outlier genes and genomic regions. Based on the finding of high variability in specific genome compartments it has been proposed that plant pathogens represent exceptional outliers in terms of evolutionary rates (Raffaele & Kamoun 2012; Upson *et al.* 2018). Nevertheless, quantitative measures of evolution in pathogen genomes are missing to test this hypothesis.

In this study we have addressed the impact of selection on genome evolution in a fungal plant pathogen, *Zymoseptoria tritici*. *Z. tritici* infects wheat and reproduces by the production of asexual spores in infected leaf tissues and by sexual recombination between isolates of opposite mating type (Waalwijk *et al.* 2002). Previous studies based on mating experiments and population genomic data have reported exceptional high recombination rates in this species (~60 cM/Mb), including intragenic recombination hotspots that underline the putative key role of recombination in evolution of this species (Croll *et al.* 2015; Stukenbrock & Dutheil 2017). The genome of *Z. tritici* consists of thirteen core and several accessory chromosomes. The latter are present at variable frequencies in different individuals, constituting a particular case of karyotypic polymorphism (Goodwin *et al.* 2011). The accessory chromosomes comprise repeat rich, heterochromatic DNA with a low gene content and they encode traits with quantitative effects on virulence (Grandaubert *et al.* 2015; Schotanus *et al.* 2015; Habig *et al.* 2017). We have previously shown that evolutionary rates on these chromosomes are particularly high suggesting that genes on the accessory chromosomes in general evolve under less selective constraints (Stukenbrock *et al.* 2010). Previous studies based on comparative population genomic analyses of *Z. tritici* and two closely related species, *Zymoseptoria pseudotritici* and *Zymoseptoria ardabiliae* used genome-wide estimates of non-synonymous and synonymous divergence to identify past species-specific signatures of selection in the wheat pathogen (Stukenbrock *et al.* 2010, 2011). Functional characterization of some of these genes revealed amino acid substitutions important for *in planta* development and asexual spore formation in the wheat-adapted pathogen, and thereby confirmed the use of evolutionary predictions to identify functionally relevant traits (Poppe *et al.* 2015).

We here apply a population genomics approach to infer genome-wide signals of natural selection, including purifying, positive and balancing selection among thirteen isolates of *Z. tritici* collected from bread wheat in Europe and the Middle East. We specifically ask to which extent recombination contributes to adaptive evolution in a sexual pathogen. Our analyses based on more than 1.4 million single nucleotide polymorphisms (SNPs) and including 700,000 coding sites allow us to identify past and on-going signatures of selection in the genome of *Z. tritici*. Our analyses reveal a strong importance of recombination in gene evolution, for both positive and negative selection, and a particularly high rate of adaptive substitutions in genes encoding putative effectors. On the other hand, balancing selection is more prevalent in genes located in repeat-rich parts of the genome implying that transposable elements also contribute to the maintenance of genetic variation. Overall, our analyses underline the potential of rapid adaptation of virulence related traits in this important agricultural pathogen.

## Results and Discussion

### Population structure of *Z. tritici* correlates with geographical origin

We generated a population genomic dataset of thirteen *Z. tritici* isolates obtained from different field populations in Europe and Iran (Table S1). Given the high extent of structural variation in genomes of *Z. tritici* isolates, we *de novo* assembled and aligned the thirteen genomes. After filtering (see Material and Methods), the resulting multiple genome alignment of ~27 Mb (Table S2) comprised a total of 1,489,362 SNPs of which approximately 50% locate in protein coding regions. The SNP data was used to compute the overall genetic diversity of the sample showing a mean value of π = 0.022 per site. Importantly, the multiple genome alignment of *de novo* assembled genomes is a priori exempt of paralogous sequences.

We first used the genomic data to investigate the relationship of the *Z. tritici* isolates. We assessed population genetic structure by analyzing the ancestral recombination graph of the thirteen genomes. To this end, we slid 10 kb windows along the multiple genome alignment, and estimated the genealogy for each window. The resulting 1,850 trees were combined into a super tree (Fig. 1A). If the sample of genomes is taken from a panmictic population with recombination, the super tree is expected to be a star tree. However, here we observe at least two clusters, one comprising all European isolates and the other comprising two isolates collected in Iran (Fig. 1A). We further investigated population structure using the program ADMIXTURE (Alexander *et al.* 2009) and found the strongest support for a model with two ancestral populations supporting the tree-based clusters of European and Iranian isolates (Fig. 1B). This pattern is consistent with some extent of geographical barriers preventing gene flow between European and Middle East *Z. tritici* populations, and possibly local adaptation to distinct host genotypes, i.e. wheat cultivars.

**Fig. 1:**
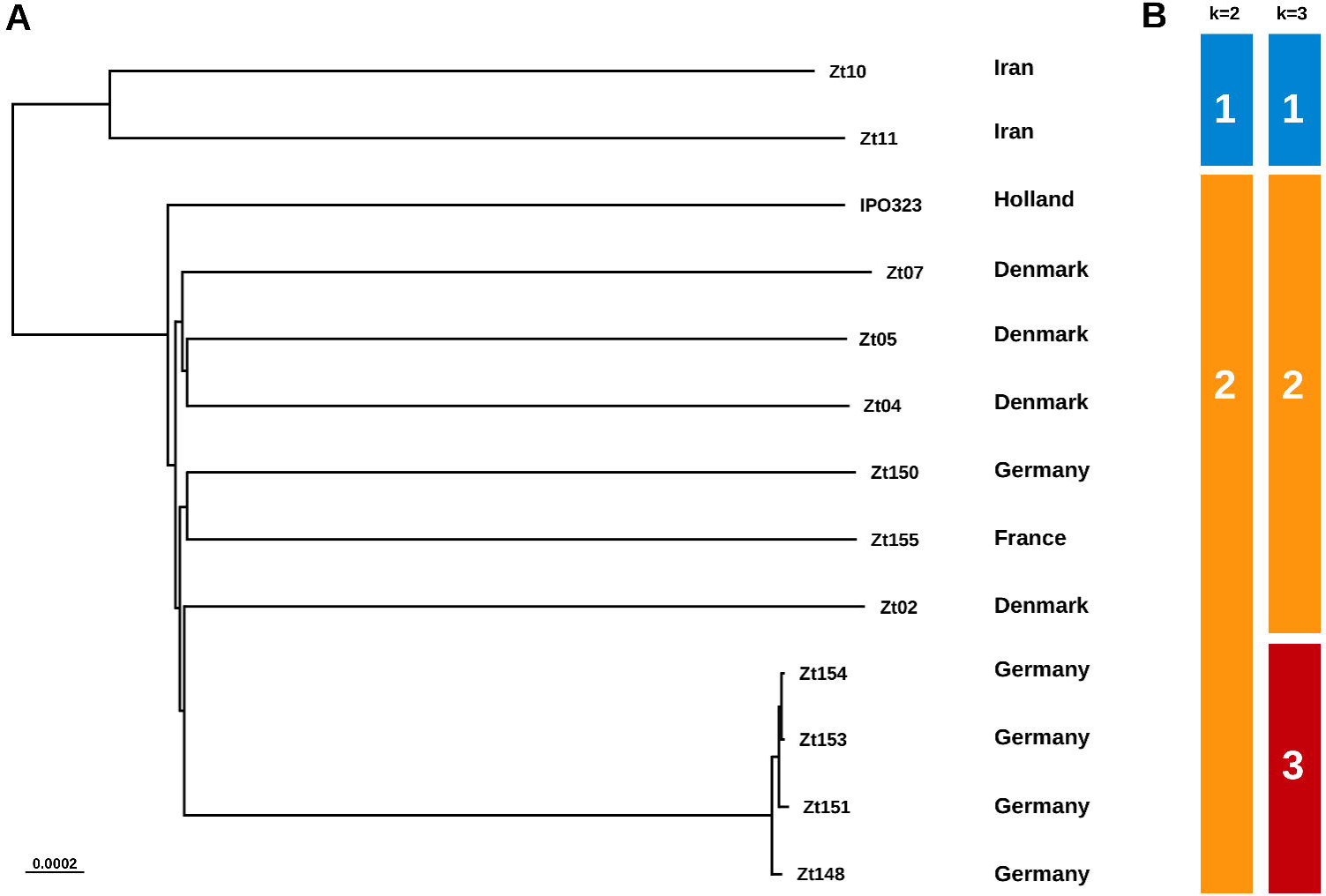
Population structure of the thirteen *Z. tritici* isolates. A) Consensus super tree of the thirteen isolates based on 1,850 genealogies estimated in 10 kb sliding windows along the multiple genome alignment. This tree suggests the grouping of the isolates into two populations originating from Europe and Iran. B) Based on SNP data, the program ADMIXTURE estimated that the best separation of the isolates is also in two populations (k=2). However, the use of k=3 highlighted a German sub-population within the European isolates.

### Recombination contributes to high rates of adaptive evolution in *Z. tritici*

We next aimed to obtain a quantitative assessment of adaptive substitutions in the genome of *Z. tritici*. To this end, we first estimated the non-synonymous and synonymous divergence dN and dS using a genome alignment of *Z. tritici* and its sister species *Z. ardabiliae*. Furthermore, we used the *Z. tritici* SNP data to compute the unfolded site frequency spectrum (SFS) of synonymous and non-synonymous sites using *Z. ardabiliae* as outgroup. The synonymous nucleotide diversity was on average over all genes 0.054, reflecting the high diversity in this species. By contrasting divergence and polymorphism data, we estimated the parameters α (proportion of adaptive non-synonymous substitutions, dN_a_/ dN) and ω_a_(proportion of the dN / dS ratio that is attributable to adaptive mutations, dN_a_/ dS). The SFS is strongly affected by demography and the presence of slightly deleterious mutations segregating at low frequencies (Eyre-Walker & Keightley 2007). State-of-the-art statistical methods account for the latter by modeling the distribution of fitness effects (DFE) of mutations (Gossmann *et al.* 2010; Galtier 2016). Potential confounding demographic factors such as variable population size, population structure and linked selection are accounted for by fitting additional parameters to accommodate deviations from a constant size neutral model of evolution. This generic correction assumes that these factors affect both synonymous and non-synonymous mutations equivalently.

We estimated α as well as ω_a_, the rate of adaptive substitutions, using four distinct DFE models accounting for mutations with both slightly deleterious and beneficial effects (see Materials and Methods) and found that the Gamma-Exponential model best fitted our data (Table 1) in agreement with studies from animals (Galtier 2016). This suggests the existence of slightly deleterious, as well as slightly beneficial segregating mutations in the genome of *Z. tritici* (Table 1). The estimates provide an α value of 35% as a genome average, and an ω_a_ value of 0.044. Both values are in the range of what is observed for Mammals (with the exception of Primates) but considerably higher than estimates from plants (Gossmann *et al.* 2010; Galtier 2016). In candidate effector genes, however, the rate of mutations fixed by selection is more than twice as high as in non-effector genes (ω_a_ equal to 0.120 vs. 0.048, Table 1), with 60% of non-synonymous substitutions in these genes inferred to be adaptive. This average estimate is close to the highest values reported in animals (Galtier 2016), and reflects the strong selective pressure acting on these genes. We note that estimates of α and ω_a_ are slightly higher when only non-effector genes are used (6,639 genes) than when using the complete gene set (6,767 genes), a small difference that likely results from sampling variance. In order to assess the significance of the observed differences between effector and non-effector genes while accounting for the difference in gene numbers in both categories (128 and 6,639 genes, respectively), we performed a bootstrap analysis where we estimated α and ω_a_ in random samples of 128 genes in each category. The results of 100 resamples are shown on Fig. 2, revealing a highly significant difference between the two distributions (Wilcoxon test, p-value < 2.2.10^−16^) and confirming the significantly higher rate of adaptation in effector genes.

**Fig. 2:**
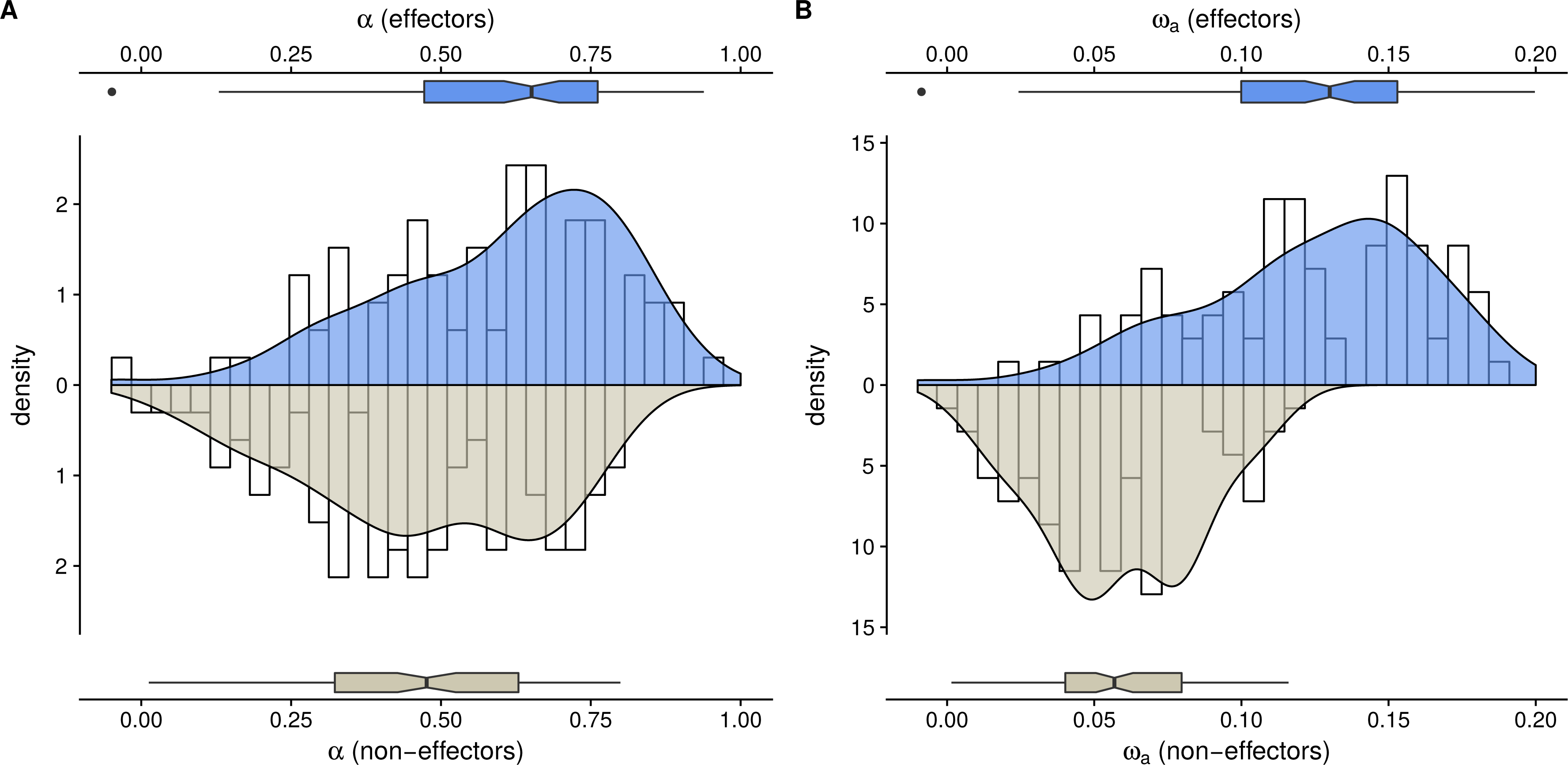
Comparison of the estimates of A) the proportion of adaptive substitution α, and B) the rate of adaptive substitution, ω_a_ for genes predicted to encode effector proteins or not. Histograms (white bars), kernel density plots and box-and-whiskers charts are computed over 100 bootstrap replicates in each case (see Material and Methods).

**Table 1:** Estimates of the proportion of adaptive mutation (α) under various models of distribution of fitness effects.

| Data set | Model | Nb. parameters | Log likelihood | AIC | $\alpha$ | $\omega_a$ |
| --- | --- | --- | --- | --- | --- | --- |
| All genes | Neutral | 16 | -1111.199 | 2254.398 | 0.253 | 0.031 |
| All genes | Gamma | 17 | -286.142 | 606.284 | 0.464 | 0.058 |
| All genes | GammaExpo | 19 | -229.425 | **496.850** | **0.352** | **0.044** |
| All genes | DisplacedGamma | 18 | -286.142 | 608.284 | 0.464 | 0.058 |
| All genes | ScaledBeta | 18 | -273.334 | 582.669 | 0.485 | 0.060 |
| All genes | BesselK | 19 | -294.387 | 626.774 | 0.514 | 0.064 |
| Non-effectors | Neutral | 16 | -1104.260 | 2240.519 | 0.252 | 0.031 |
| Non-effectors | Gamma | 17 | -286.821 | 607.643 | 0.463 | 0.057 |
| Non-effectors | GammaExpo | 19 | -230.221 | **498.442** | **0.388** | **0.048** |
| Non-effectors | DisplacedGamma | 18 | -286.821 | 609.643 | 0.463 | 0.057 |
| Non-effectors | ScaledBeta | 18 | -273.687 | 583.375 | 0.458 | 0.057 |
| Non-effectors | BesselK | 19 | -296.728 | 631.456 | 0.513 | 0.063 |
| Effectors | Neutral | 16 | -101.036 | 234.071 | 0.307 | 0.062 |
| Effectors | Gamma | 17 | -94.255 | 222.510 | 0.485 | 0.097 |
| Effectors | GammaExpo | 19 | -87.332 | 212.664 | 0.666 | 0.134 |
| Effectors | DisplacedGamma | 18 | -94.684 | 225.369 | 0.492 | 0.099 |
| Effectors | ScaledBeta | 18 | -86.845 | **209.689** | **0.600** | **0.120** |
| Effectors | BesselK | 19 | -87.416 | 212.832 | 0.507 | 0.102 |

AIC: Akaike's information criterion. $\alpha$ : proportion of adaptive substitutions, $\omega_a$ : rate of adaptive substitutions. Values in bold indicate the best model fit for each gene set.

**Table 2:**
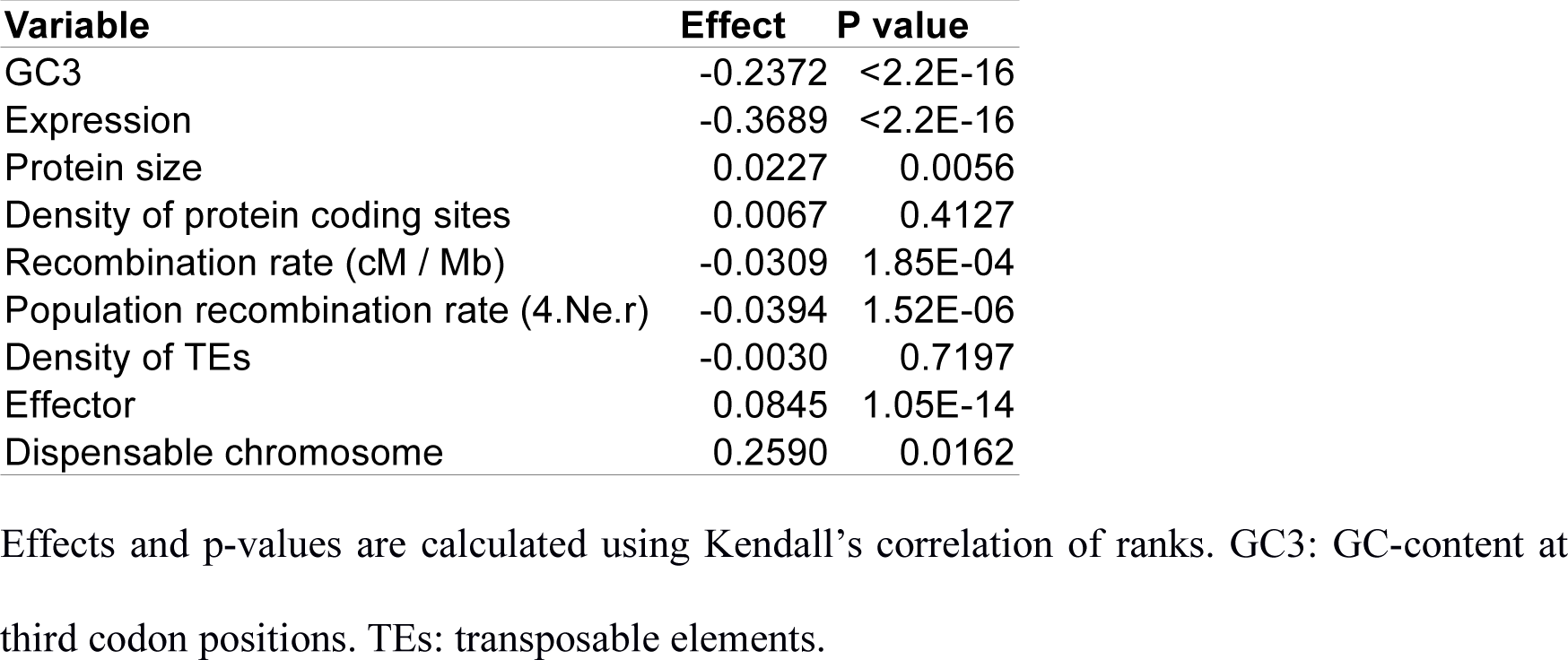
Correlation of pN / pS with genomic factors.

We hypothesized that recombination could be an important driver of adaptive evolution in *Z. tritici*. Previous inference of recombination maps in *Z. tritici* based on experimental crosses and population genomic data have revealed exceptionally high rates of recombination in this species (Croll *et al.* 2015; Stukenbrock & Dutheil 2017). To assess the role of recombination in adaptive evolution of *Z. tritici* we used the recombination maps generated in these previous studies. Genetic maps resulting from crossing experiments allow inference of the recombination rate r (measured as cM / Mb), but are limited in resolution. Conversely, linkage disequilibrium-based maps generated from population genomic data offer an improved resolution, but only allow inference of ρ = 4.Ne.r, where Ne is the effective population size. As such, ρ is a proxy for r that is affected by both selection and demography. We clustered all analyzed genes according to their r and ρ values, and estimated α and ω_a_ for each case using the Gamma-Exponential distribution of fitness effects. In order to assess the variance of our estimates and their robustness to the sampled genes, we further conducted a bootstrap analysis where we sampled genes in each category 100 times. We report a significant positive correlation between α (averaged over 100 bootstrap replicates) and r (Kendall’s tau = 0.31, p-value = 0.004354) and ω_a_ and r (Kendall’s tau = 0.31, p-value = 0.006041). We note that similar correlations are observed when ρ is used instead of r, or when effector genes are discarded (Supplementary Data). These results suggest that a higher recombination rate favors the fixation of adaptive mutations, as expected under a Hill-Robertson interference scenario, where selected mutations reduce the effective population size at linked loci (Hill & Robertson 1966; Marais & Charlesworth 2003).

We further explored the relationship between α, ω_a_ and r. We fitted four models: linear (as in (Campos *et al.* 2014)), power law, curvilinear (as in (Castellano *et al.* 2016)), and logarithmic (see Materials and Methods). While we find a higher support for the logarithmic model (Fig. 3), the effect is very weak and our data does not allow further estimation of the asymptotic value (Castellano *et al.* 2016). When using ρ instead of r, the curvilinear model is preferred when all genes are considered, but the power law offers a better fit when effectors are excluded (Supplementary Material).

**Fig. 3:**
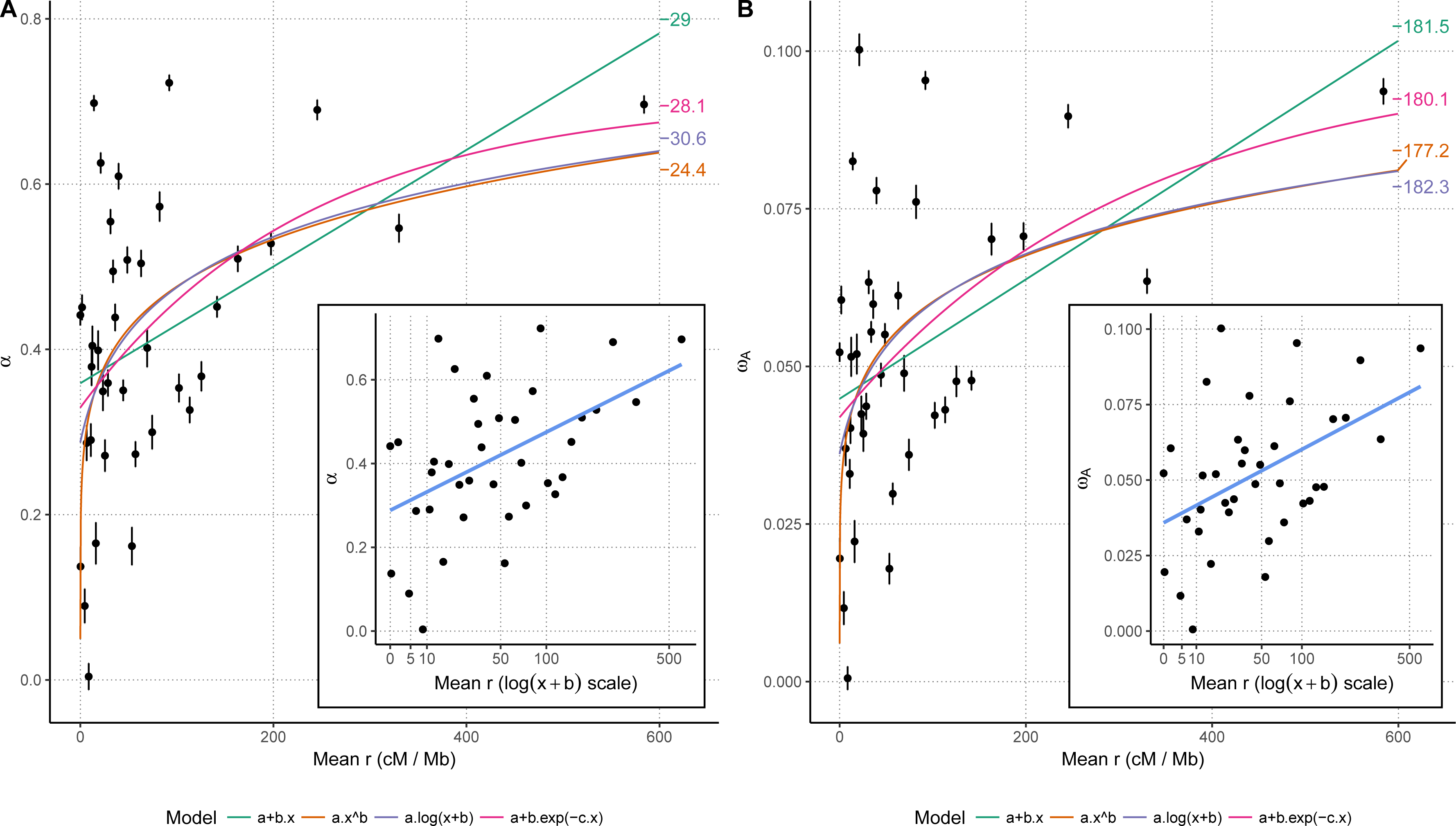
Estimates of A) the proportion of adaptive substitution α, and B) the rate of adaptive substitution, ω_a_ as a function of the recombination rate (r). Each point and bars represent the mean estimate and corresponding standard error for one recombination category over 100 bootstrap replicates. Four models were fitted (colored curved) and corresponding Akaike’s information criterion values are indicated in the right margin. Inset plots represent the same data with a logarithmic scale, the b value was set to the corresponding estimate in the third model. Confidence intervals have been omitted for clarity.

In summary, our results represent a quantitative assessment of adaptive evolution in the genome of a fungal pathogen and reveal a strong role of recombination on adaptation (Fig. 3). The exceptionally high rate of adaptation in effector genes (Fig. 2) likely reflects arms race evolution driven by the antagonistic interaction of *Z. tritici* and its host.

### Local rates of recombination are correlated with the strength of purifying selection revealing pervasive background selection

We next addressed the genome-wide strength of purifying selection in protein coding genes of *Z. tritici* using the ratio of non-synonymous to synonymous polymorphisms (pN / pS ratio). We computed pN and pS for each gene as the average pairwise heterozygosity (Romiguier *et al.* 2014; Ellegren & Galtier 2016). To investigate which genome parameters impact the strength of purifying selection in *Z. tritici*, we compared the pN / pS ratio for each gene to 1) the mean gene expression, 2) the GC content at third codon positions (GC3), 3) the protein length, 4) the local recombination rate, 5) the density in protein coding sites and 6) the density in transposable elements. We used *Z. tritici* gene expression data from early host colonization (four days after spore inoculation on leaves of seedlings of a susceptible wheat host) and *in vitro* growth (Kellner *et al.* 2014). Recombination rates were averaged in 20 kb windows and recombination was analysed independently as r (Croll *et al.* 2015) and ρ (Stukenbrock & Dutheil 2017). We further considered whether the gene 7) is an effector candidate and 8) is located on an accessory chromosome (Fig. 4). We restricted our analysis to genes for which pN / pS could be computed (6,627 genes, see Materials and Methods) and for which pN / pS was estimated to be < 1 (6,621 genes).

**Fig. 4:**
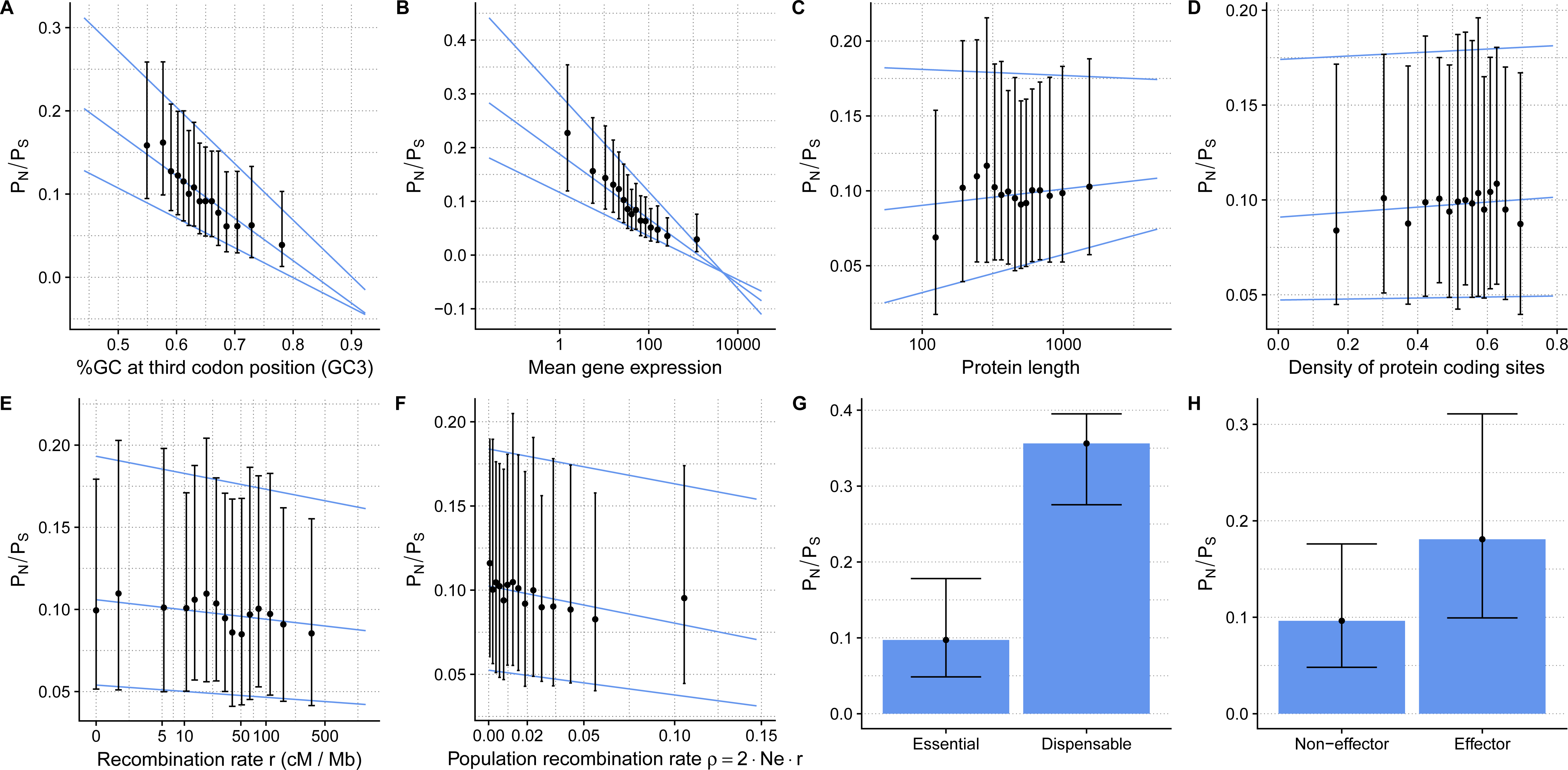
Correlation of the strength of purifying selection with several genomic factors. The intensity of purifying selection is measured by the pN / pS ratio. Points represent median values and error bar the first and third quartiles of the distributions. A-F: x-axis were discretized in categories with equal point densities for clarity of visualization. Lines represent first, median and third quantile regression on non-discretized data.

We identify several variables that significantly impact the strength of purifying selection (summarized in Table 2). Mean gene expression and GC at third codon position have the strongest effect on pN / pS (Fig. 4A and 4B), displaying highly significant negative correlations (Kendall’s tau = −0.369 and −0.237 respectively, p-values < 2.2.10^−16^ in both cases). Consistent with this observation, studies in yeast and bacteria have previously documented a strong impact of expression levels on gene evolution whereby highly expressed genes are more conserved reflected as lower pN / pS values (Drummond *et al.* 2006; Liao *et al.* 2006). GC3 and mean gene expression are intrinsically highly correlated (Kendall’s tau = 0.222, p-value < 2.2.10^−16^), possibly reflecting biases in codon usage whereby optimal codons are GC-rich at their third position (Fig. S1), as also observed is other organisms (Duret & Mouchiroud 1999). An alternative explanation for the effect of the GC content on pN / pS could be a possible indirect effect of recombination as we also observe a positive correlation of GC3 and the recombination rate (Kendall’s tau = 0.097, p-value < 2.2.10^−16^). A similar correlation of recombination and GC3 is found in other organisms (Duret 2002). In *Saccharomyces cerevisiae* this correlation has been explained by the impact of biased gene conversion on sequences evolution (Birdsell 2002). However, a thorough search for signatures of GC-biased gene conversion did not find any pervasive effect of this phenomenon in *Z. tritici* (Stukenbrock & Dutheil 2017). The relationship between pN / pS and GC3 is therefore more likely a by-product of the correlation with gene expression.

Protein size is slightly positively correlated with pN / pS (Table 2), although the effect is due to very short proteins being more conserved and the observed effect perishes when testing only proteins with > 100 amino acids (excluding 348 proteins out of 6,621, Kendall’s tau = 0.012, p-value = 0.1443, Fig. 4C). Gene density, estimated in a 50 kb regions centered on the gene (see Material and Methods), does not have a significant effect on pN / pS (Kendall’s tau = 0.0067, p-value = 0.4127, Fig. 4D). The genome of *Z. tritici* is compact and uniform in terms of gene localization, and the distribution of gene density is almost normal with a median around 54%. This likely explains that gene density does not have an impact on strength of purifying selection. Conversely, we observe a significant negative correlation between pN / pS and recombination rate r (Kendall’s tau = −0.031, p-value = 1.85.10^−4^, Fig. 4E) or ρ (Kendall’s tau = −0.039, p-value = 1.52.10^−6^, Fig. 4F and Fig. 5). These results are in agreement with a model of background selection, where purifying selection at linked loci with low recombination rates reduces the local effective population size, therefore reducing the efficacy of selection and allowing slightly deleterious mutations to spread more frequently than at loci with high recombination rates (Charlesworth *et al.* 1993; Nordborg *et al.* 1996).

**Fig. 5:**
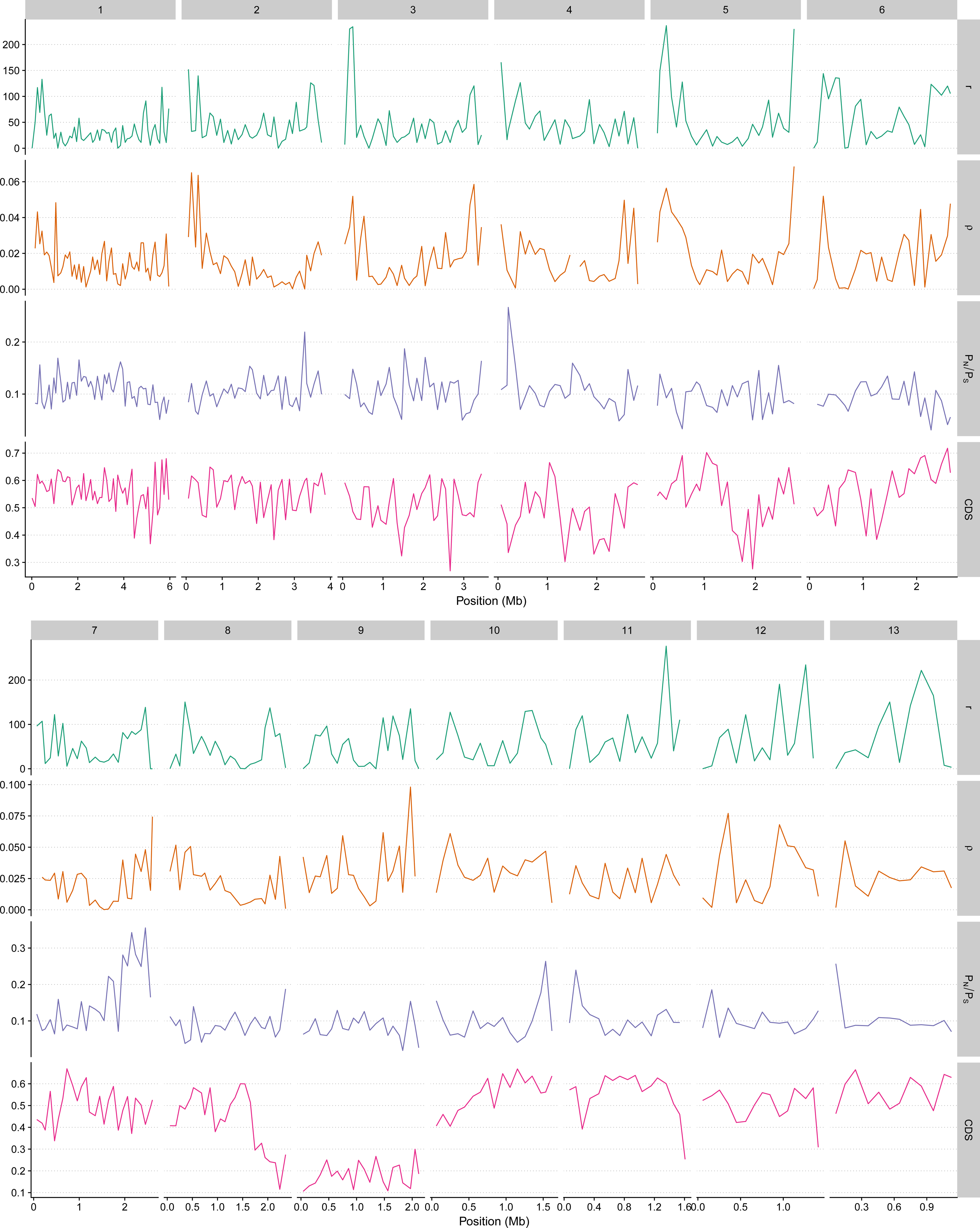
Patterns of selection along the genome of *Z. tritici*. Recombination rate, population recombination rate, pN / pS ratio and density of coding sites (CDS) are plotted in windows of 100 kb along the thirteen essential chromosomes.

Background selection is notably expected to be stronger in regions of higher density of coding sites, implying that the negative correlation between recombination and pN / pS should be higher in gene-dense regions. To further test this effect of coding site density, we split our gene set in two subsets, whether the density of coding sites was below (low-density set) or above (high-density set) the median. For the low-density set, we report a marginally significant negative correlation between r and pN / pS (Kendall’s tau = −0.022, p-value = 0.05856), while the correlation is stronger and significant for the high-density set (Kendall’s tau = −0.039, p-value = 7.89.10^−04^). These results provide evidence that background selection impacts the genome of *Z. tritici*, and support a central role of recombination in the removal of non-adaptive mutations in the genome of *Z. tritici* consistent with patterns described in other species such as *Drosophila melanogaster* (Campos *et al.* 2014). We note, however, that the effect of recombination is smaller than the one of functional variables such as mean expression. This is likely related, we hypothesize, to the globally high level of recombination and reduced linkage throughout the genome of *Z. tritici*.

One part of the *Z. tritici* genome where recombination is low is the accessory chromosomes (Stukenbrock & Dutheil 2017). This low recombination rate is reflected in our estimates of purifying selection. We find a significantly higher pN / pS ratios in genes located on accessory chromosomes (Wilcoxon rank test, p-value = 0.0162, Fig. 4G), and on the right arm of chromosome 7 (Fig. 5), a genomic region predicted to be an ancestral accessory chromosome fused with a core chromosome (Schotanus *et al.* 2015). Accessory chromosomes have a reduced effective population size due to their presence/absence variation among individuals, resulting in a reduced efficacy of selection and a higher pN / pS ratio on these chromosomes..

Finally, we compared the strength of purifying selection of effector and non-effector genes, and find that genes predicted to encode effector proteins have a significantly higher pN / pS ratio compared to other genes (Wilcoxon rank test, p-value = 1.5.10^−14^, Fig. 4H). We speculate that this pattern is due to the fast evolution through positive selection of this particular category of genes, and the higher pN / pS ratio in these genes reflects the fixation of slightly deleterious mutations by linkage.

### Detection of on-going balancing selection in *Z. tritici* identifies candidate pathogenicity factors

The recurrent interaction with different host genotypes can confer the maintenance of multiple alleles at selected loci in the pathogen population. To identify specific sites and genes in the *Z. tritici* genome showing signatures of balancing selection, we fitted models of codon sequence evolution as implemented in the CodeML program of the PAML package to detect genes with significant signatures of balancing selection using likelihood ratio tests (Yang 2007). Two models are typically compared: a model with sites evolving only under neutrality or purifying selection, with an ω ratio (non-synonymous rate of polymorphisms / synonymous rate of polymorphisms) equal or below one (neutral model with purifying selection), and a model allowing for some sites to evolve under positive selection with a ω ratio above one (positive selection model). A likelihood ratio test (LRT) is then used to test for the occurrence of positive selection. When applied to population data, sites with ω > 1 can reflect balancing selection (Anisimova *et al.* 2001). We fitted codon models for genes present in at least three isolates (83% of genes located on core chromosomes and 31% of genes on the accessory chromosomes, Table S3). After correcting for multiple testing, we identified a final set of 787 genes (including 24 on the accessory chromosomes) evolving under balancing selection (false discovery rate < 0.01, Table S3).

As selection tests based on codon model comparison were previously shown to potentially suffer from an inflated false discovery rate (FDR) in the presence of recombination (Anisimova *et al.* 2003), we conducted simulations with parameters reflecting the characteristics of our data set (see Material and Methods). In agreement with previous results, we report an increase in FDR in the presence of recombination within the gene (Fig. 6). While our results appear to be relatively independent of the level of diversity in the alignment, we report a strong effect of the number of sites in the alignment: for a given number of recombination events, we see a higher FDR in long compared to short genes. This suggests that the recombination rate, which is lower in longer genes for a given number of recombination events, is not the only determinant of erroneous rejection of the null model. Large alignments, on the contrary, carry more statistical signal (Anisimova *et al.* 2001) and can lead to strong support of the wrong model in case an incorrect tree is provided, an effect that is independent of the actual number of recombination events. In agreement with this hypothesis, we see that the FDR also increases with the number of sequences in the alignment (Fig. 6). To assess the extent of false discovery in our analysis, we sorted genes for which all 13 individuals were present (6,627 genes) according to their corresponding protein lengths: more than 100, 500, 1,000 and 2,000 amino acids, respectively. We find that the proportion of genes significantly rejecting the null hypothesis of no positive selection is systematically higher than the maximum observed FDR for the corresponding length (Fig. 6). This suggests that false discovery due to recombination does not explain all of our candidates and that the selection test was able to capture biological signal.

**Fig. 6:**
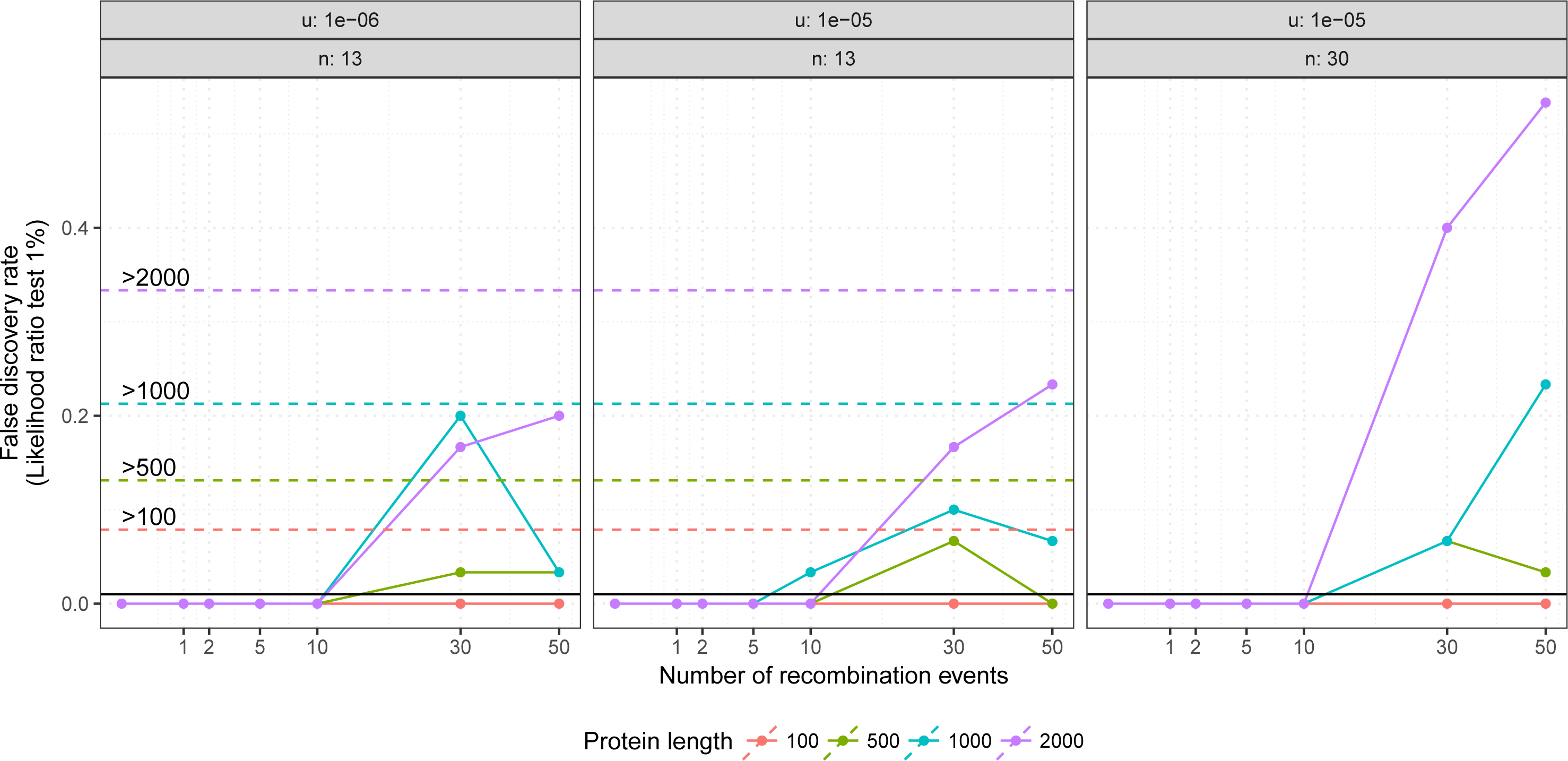
Effect of recombination on the inference of positive selection. False discovery rate, as estimated from simulations under a model with neutral and purifying selection only, was plotted as a function of the number of recombination events (x-axis), length of the alignment (coloured lines), mutation rate and sample size (panels). Horizontal dash lines show the discovery rate of the real data for distinct minimum gene lengths.

To further assess the gene-specific selection regime, we computed Tajima’s D for each gene in our data set. We find that the distribution of Tajima’s D is globally shifted toward negative values (Fig. 7), as expected for coding sequences under purifying selection. Furthermore, *Z. tritici* was previously found to have undergone population expansion since the domestication of wheat and speciation of the pathogen (Stukenbrock *et al.* 2007). This population expansion also contributed to the overall negative Tajima’s D values. We further observe that there is no significant differences between genes encoding predicted effector proteins and others (Wilcoxon rank test, p-value = 0.1597). Genes predicted to be under balancing selection by PAML, however, display significantly higher Tajimas’ D values (Wilcoxon rank test, p-value < 2.2.10^−16^), a typical signature of balancing selection (Tajima 1989). However, given the population structure that we observe, it cannot be excluded that for some of these genes, the signal of the LRT results from population differentiation. In such case, the detected gene could be under positive selection resulting from local adaptation. In the following, we further investigate the genome distribution and biological function of genes detected by PAML.

**Fig. 7:**
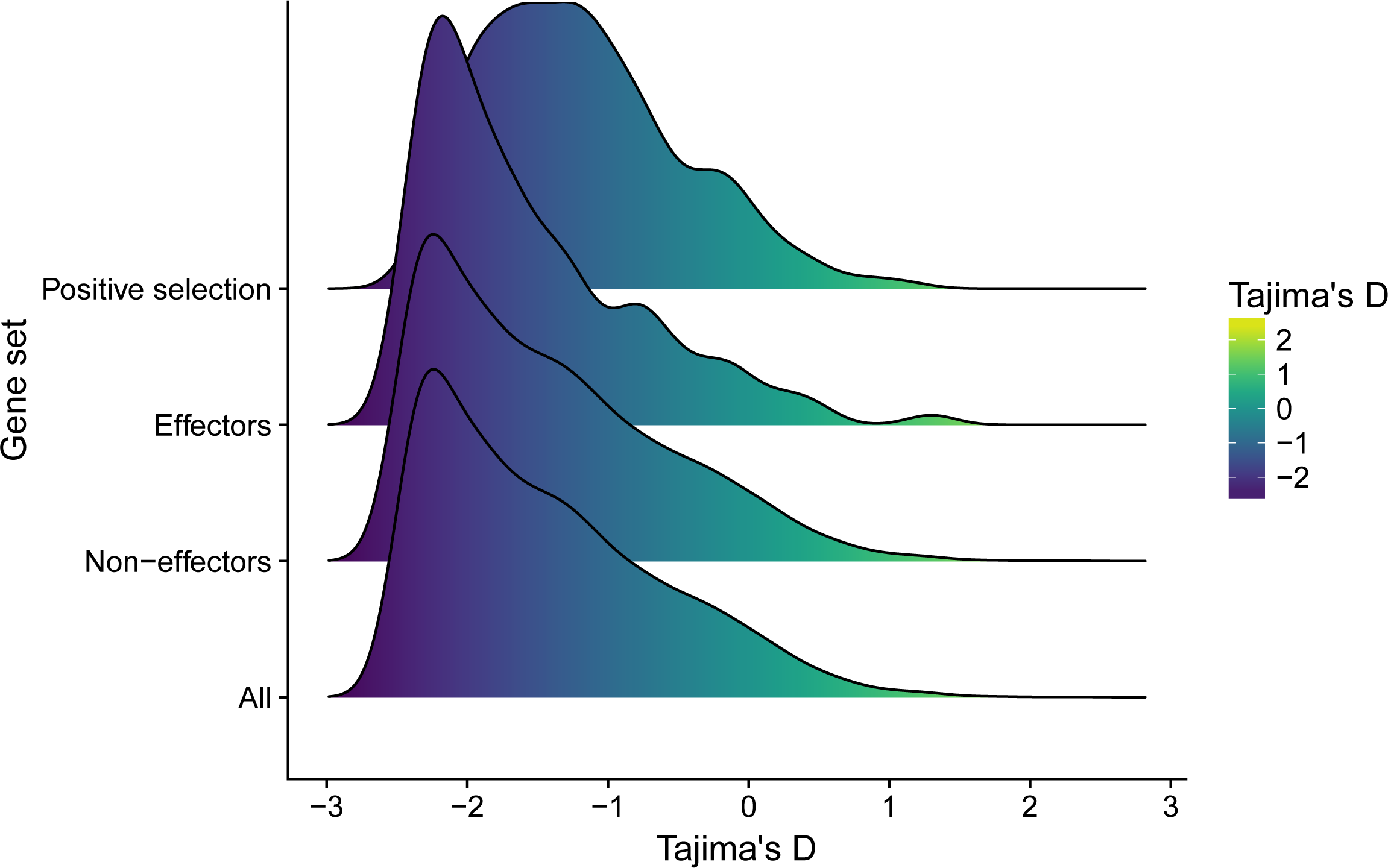
Distribution of Tajima’s D for different gene categories. Kernel densities were fitted to the distribution of each gene’s Tajima’s D (x-axis and color scale), sorted per category (detected to be under balancing selection, predicted to encode an effector protein, predicted not to encode an effector protein, all genes).

In several pathogen genomes, rapidly evolving genes are found clustered in particular genomic environments often associated with repetitive sequences (e.g (Raffaele *et al.* 2010; Dutheil *et al.* 2016)). To address whether the same pattern is found in *Z. tritici*, we assessed the spatial distribution of genes under balancing selection along chromosomes (see Materials and Methods). Of the 787 positively selected genes, 240 are located within a distance of 5 kb from each other and thereby form 108 clusters containing two to four genes with signatures of positive selection. At the genome scale, clusters containing two or three genes do not show a significant pattern as the same pattern can be obtained by randomly distributing the positively selected genes across the genome. However, there are two significant clusters (p-value = 1.8.10^−3^) containing four and eight genes, respectively. The clusters comprising four genes is located in a 24 kb region of chromosome 5, and the cluster comprising eight genes in a 31 kb region of chromosome 9. For genes in both clusters no functional relevance can be assigned, but the clusters represent interesting candidates for future functional studies.

### The genomic determinants of balancing selection in *Z. tritici*

In order to test which factor drives the occurrence of balancing selection in the *Z. tritici* genome, we fitted (generalized) linear models. We assessed the impact of 1) the recombination rate, 2) the density in protein coding sites, 3) the density in transposable elements (TE), 4) whether the gene is predicted to encode an effector protein, and 5) the mean gene expression (see Material and Methods). We fitted a binary logistic model, where the response variable is whether a gene is predicted to be under positive balancing selection by PAML. We find that positive selection is less likely at highly expressed genes (Table 3), an effect that, we hypothesize, is due to highly expressed genes being on average more constrained (see above).

**Table 3:**
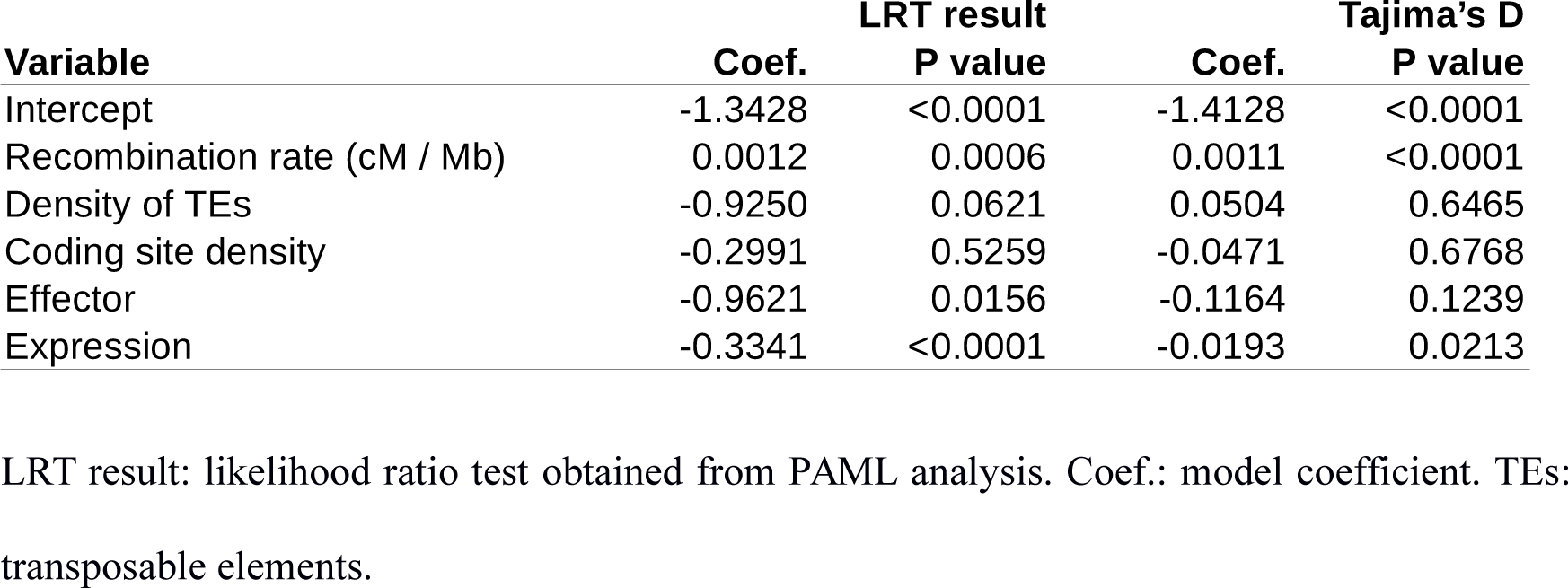
Genomic factors affecting the occurrence of balancing selection

We find a significant effect of effector encoding genes (Table 3), that is, effector genes are, intriguingly, less likely to be under balancing selection than non-effector encoding genes. As described above, we find that effector genes tend to undergo a higher rate of adaptive evolution, and our analyses thereby suggest that distinct categories of genes are evolving under arms race (recurrent selective sweeps) and trench-warfare (balancing selection) scenarios in the pathogen genome. The rate of recombination has a weak but significant positive effect on the occurrence of positive selection, *i.e.* genes detected to be under balancing selection are more frequent in highly recombining regions (Table 3). This effect can be interpreted as a better efficacy of selection in highly recombining regions, but can also be due to an increased false discovery rate in the presence of recombination. In order to disentangle the two hypotheses, we fitted a similar linear model with Tajima’s D as a response variable and find consistent results where recombination rate has a significant positive effect on Tajima’s D (Table 3).

We further extended our analyses of the genes predicted to be under balancing selection by characterizing known protein domains. From the 787 candidate genes, 602 of the encoded proteins (76.5%) have an *in silico* attributed function or harbor known protein domains. We conducted a PFAM domain enrichment analyses and identified 21 significantly enriched domains (FDR <= 0.05) (Table S4). These domains can be grouped into different categories based on their associated molecular function and include carbohydrate-active enzymes (CAZymes), polyketide synthases (PKS), non-ribosomal peptide synthetases (NRPS), and cellular transporters (Table S4). Several of these categories are relevant for the pathogen to interact with its host. For example, CAZymes are proteins involved in the break down of glycosidic bonds contained in plant cell walls (André *et al.* 2014), and PKS and NRPS are multimodular enzymes involved in the biosynthesis of secondary metabolites of which many also have been shown to be involved in virulence of plant pathogenic fungi (Howlett 2006). Among the positively selected *Z. tritici* genes, we also searched for homologs of known virulence factors described in other plant pathogens. The PFAM domain PF14856 corresponds to the mature part of a virulence factor Ecp2 described in the tomato fungal pathogen *Cladosporium fulvum* (Van den Ackerveken *et al.* 1993). Ecp2 has been described in several other plant pathogens (Stergiopoulos *et al.* 2010) and has three homologs in *Z. tritici*. We find that two of these homologs comprise sites under positive selection supporting a virulence related role of this gene also in this wheat pathogen.

### Signatures of past selection in *Z. tritici* and related species

In a previous study the evolutionary history of *Z. tritici* was inferred using a whole genome coalescence analyses (Stukenbrock *et al.* 2011). We showed that divergence of *Z. tritici* and its sister species *Z. pseudotritici* occurred recently and likely coincides with the onset of wheat domestication and thereby specialization of *Z. tritici* to a new host. We hypothesize that genes important for the colonization of distinct hosts have been under selection during the divergence of *Zymoseptori*a species. To infer signatures of past selection we applied the branch model implemented in the program package PAML to estimate the branch-specific dN / dS ratio for core *Zymoseptoria* genes (present in four species *Z. tritici*, *Z. pseudotritici*, *Z. ardabiliae* and *Z. brevis*) (Yang & Nielsen 1998; Grandaubert *et al.* 2015). The branch-specific dN / dS ratios reflect the proportion of non-synonymous to synonymous substitutions accumulated in each branch of the *Zymoseptoria* phylogeny. Our analyses identified 47 genes with a dN /dS ratio > 1 on the *Z. tritici* branch indicative of an increased non-synonymous divergence, 54 genes in *Z. pseudotritici*, 60 genes in *Z. brevis* and 15 genes in *Z. ardabiliae* (Table 4). Based on their putative function in host-pathogen interactions, we hypothesized that some positively selected genes encode secreted proteins and putative effectors. In order to test this hypothesis, we fitted linear models with branch specific dN / dS ratios as response variables and whether the corresponding gene family encodes an effector or not in *Z. tritici* as explanatory variable. For all four extant species and the common ancestor of *Z. tritici* and *Z. pseudotritici*, we find a significantly higher dN / dS ratio for genes encoding predicted effector proteins (Table 4), in agreement with the general observation that effector-encoding genes are fast-evolving. For *Z. tritici* only we report that effector-encoding genes are more likely to have a dN / dS > 1. For *Z. ardabiliae*, we only find 15 genes under positive selection, and none in effector-encoding candidate genes.

**Table 4:**
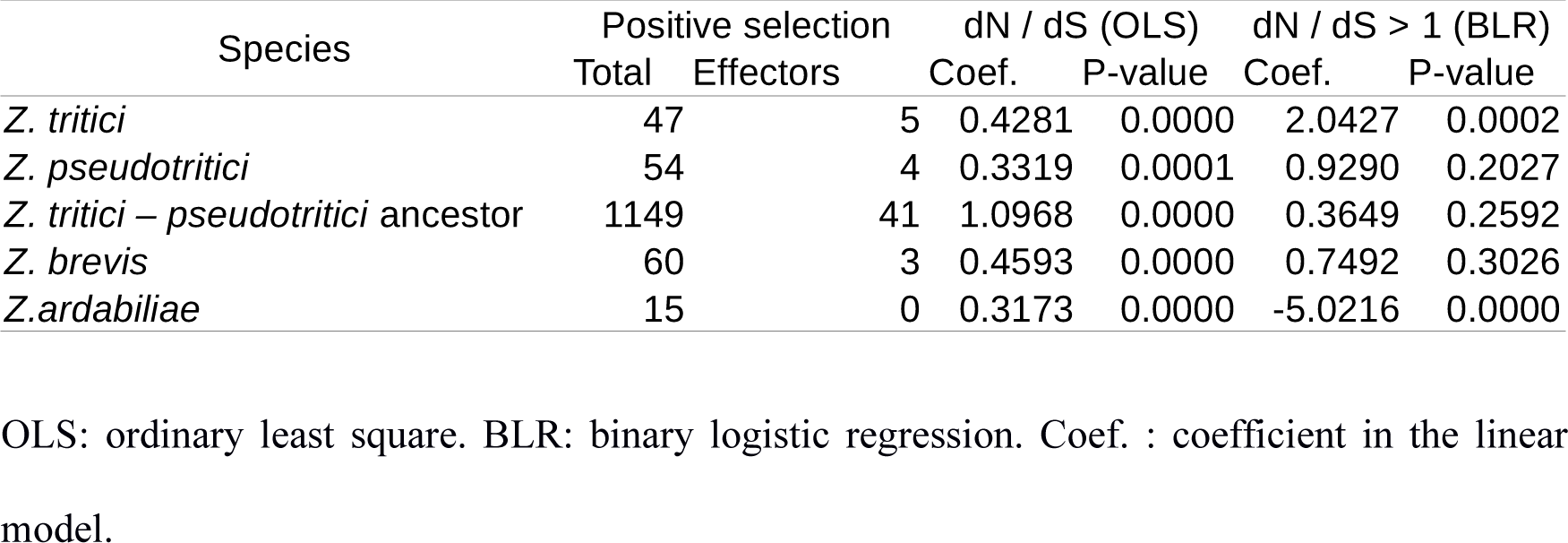
Difference of dN / dS ratios between genes predicted to encode an effector protein or not, in different branches of the species tree.

We next performed a PFAM domain enrichment analysis for the positively selected genes with a predicted function to address if some functional domains are significantly enriched in this set of genes (FDR < 0.05) (Table S5). The majority of genes encode proteins of unknown function, however among the functionally characterized proteins we find a gene encoding a regulator of chromosome condensation that was previously described to be functionally relevant for virulence in *Z. tritici* (Poppe *et al.* 2015). Our analyses reveal an enrichment of genes encoding proteinases, one gene encoding Lysin motifs (LysM) already described as an effector in *Z. tritici* and other fungal pathogens (de Jonge *et al.* 2010; Marshall *et al.* 2011), and one gene encoding a CFEM domain. Cystein-rich CFEM domains have been described in G-protein-coupled receptors in the rice blast pathogen *Magnaporthe oryzae* playing an important role in pathogenicity (Kulkarni *et al.* 2005). These genes provide interesting candidates for further studies of molecular determinants of host specificity in *Z. tritici*.

## Conclusions

In this study we have used the fungal wheat pathogen *Z. tritici* to assess the patterns of selection acting on the genome, both qualitatively and quantitatively. We measure the rate of adaptation using models of distributions of fitness effects, and with polymorphism and divergence models of codon sequence evolution, we provide evidence for signatures of both positive and balancing selection in protein coding genes, as expected under arms race and trench warfare scenarios, respectively. The rate of adaptive substitutions in the plant pathogen is similar to estimates in animal species and considerably faster than corresponding estimates in plants. Notably, rates of evolution in genes encoding effector proteins are more than twice as fast as the genome average. Furthermore, our results suggest widespread occurrence of linked selection (both Hill-Robertson interference and background selection), as both the rate of adaptation and the strength of negative selection correlate with the recombination rate. Finally, we infer signatures of balancing selection and find an enrichment of genes encoding pathogenicity related functions - but not effector proteins - among detected genes. Our results thereby demonstrate that different categories of genes evolve under arms race and trench-warfare selection in this pathogen. Interestingly, we show that signatures of positive selection and balancing selection do not correlate with the presence of transposable elements as predicted in other studies. These results highlight the fundamental role of recombination and sexual reproduction in adaptive processes of rapidly evolving organisms.

## Materials and Methods

### Re-sequencing, assembly and alignment of Z. tritici isolates

In this study we used a geographical collection of thirteen field isolates of *Z. tritici* isolated from infected leaves of bread wheat (*Triticum aestivum*) (Table S1). Genome data of three isolates, including the reference isolate IPO323, were published previously (Goodwin *et al.* 2011; Stukenbrock *et al.* 2011). For the remaining ten isolates full genomes were sequenced. DNA extraction was performed as previously described (Stukenbrock *et al.* 2011). Library preparation and paired end sequencing using an Illumina HiSeq2000 platform were conducted at Aros, Skejby, Denmark. Sequence data of the ten isolates has been deposited under the NCBI BioProject IDs PRJNA312067. We used SOAPdenovo2 (Luo *et al.* 2012) to construct de novo genome assemblies for each isolate independently. For each genome, the k-mer value maximizing the weighted median (N50) of contigs and scaffolds was selected.

Prior to generating a multiple genome alignment, we pre-processed the individual genomes of the thirteen *Z. tritici* isolates. First, we masked repetitive sequences using a library of 497 repeat families identified de novo in four *Zymoseptoria* species (Grandaubert *et al.* 2015). Repeats were soft-masked using the program RepeatMasker (option -xsmall) to retained information of repeat sites in the alignment (A.F.A. Smit, R. Hubley & P. Green RepeatMasker at http://repeatmasker.org). Second, we filtered the genome assemblies to contain only contigs with a length >= 1 kb. Multiple genome alignments were generated by the MULTIZ program using the LASTZ pairwise aligner from the Threaded Blockset Aligner (TBA) package (Blanchette *et al.* 2004). The alignment was projected on the IPO323 reference genome using the maf_project program from the TBA package.

### Inference of population structure

In order to infer population structure, we generated genealogies of the thirteen isolates using the multiple genome alignment. We used the MafFilter program (Dutheil *et al.* 2014) to compute pairwise distance matrices using maximum likelihood under a Kimura 2 parameter model in 10 kb sliding windows along the chromosomes of the reference genome. For each window, a BioNJ tree was reconstructed from the distance matrices. The resulting 1,850 genealogies were used to build a super tree using SDM for the generation of a distance supermatrix (Criscuolo *et al.* 2006) and FastME was used to infer a consensus tree (Lefort *et al.* 2015). We used the program ADMIXTURE (Alexander *et al.* 2009) (software using the — haploid=’*’ option) to estimate the number of ancestral populations based on single nucleotide polymorphism data. Filtered biallelic variants were exported as PLINK files using MafFilter (Dutheil *et al.* 2014). A cross-validation procedure was conducted as described in the manual of the ADMIXTURE in order to determine the optimal number of partitions. We further assessed the effect of linkage by removing SNPs with an R2 value higher than 0.1 with any other SNP in windows of 50 SNPs, slid by 10 SNPs, using PLINK (Chang *et al.* 2015). This filtering did not affect the conclusion of the cross-validation procedure.

### Prediction of effector candidates

Gene models from the *Z. tritici* reference strains (Grandaubert *et al.* 2015) were used to predict proteins targeted for secretion using SignalP (Petersen *et al.* 2011). Genes predicted to encode a secreted protein were further submitted to effector prediction using the EffectorP software (Sperschneider *et al.* 2016).

### Estimation of rates of adaptation

Based on the coordinates of each predicted gene model in the reference genome IPO323 (Goodwin *et al.* 2011; Grandaubert *et al.* 2015), exons were extracted from the multiple genome alignment of *Z. tritici* isolates using MafFilter (Dutheil *et al.* 2014). Complete coding sequences (CDS) were concatenated to generate individual alignments of all orthologous CDS. If one or more exons were not extracted from the alignment due to missing information, the gene was discarded from further analyses. Each complete CDS alignment was filtered according to the following criteria: (i) CDS were discarded if they contained more than 5% gaps in one or more individuals, (ii) CDS with premature stop codon were likewise deleted, and (iii) only alignments comprising three or more CDS were kept. In some cases, due to indels in the genome alignment, the codon phasing of some genes was lost. This issue was overcome by refining the CDS alignment using the codon-based multiple alignment program MACSE (Ranwez *et al.* 2011). The final data set contains 9,412 gene alignments, among which 7,040 contain a sequence for all 13 isolates. We further created a data set containing an outgroup sequence, taken from *Z. ardabiliae*, leading to 6,767 alignments with all 13 isolates together with the outgroup sequence.

The CDS alignment with outgroup was used to infer the synonymous and non-synonymous divergence based on the rate of synonymous and non-synonymous substitutions. The synonymous and non-synonymous unfolded site frequency spectra (SFS) were computed, using the outgroup sequence to reconstruct the ancestral allele. To do so, we first reconstructed a BioNJ tree for each gene and fitted a codon model of evolution using maximum likelihood. Then ancestral state was inferred using the marginal reconstruction procedure of Yang (Yang *et al.* 1995). All calculations were performed using the BppPopStats program from the Bio++ Program Suite (Guéguen *et al.* 2013). We used the Grapes program in order to estimate the distribution of fitness effects from the SFS and compute a genome wide estimate of α and ω_a_, the proportion of mutations fixed by selection and the rate of adaptive substitutions respectively (Galtier 2016). The following models were fitted and compared using Akaike’s information criterion: Neutral, Gamma, Gamma-Exponential, Displaced Gamma, Scaled Beta and Bessel K. Analyses were conducted on the complete set of gene alignments, as well as on sub-datasets sorted according to whether the individual genes encoded a predicted effector protein or not. We further stratified our data set according to the local recombination rate, computed in 20 kb windows, using both the previously published genetic maps (Croll *et al.* 2015) and population estimates from patterns of linkage disequilibrium (Stukenbrock & Dutheil 2017). We discretized the observed distributions of both r and ρ in 41 and 45 categories, respectively, using the cut2 command from the Hmisc R package in order to have similar number of genes in each category (comprising between 247 and 258 genes for ρ, and between 67 and 1,323 genes for r, the largest value being obtained for genes with r = 0). For each gene sets, 100 bootstrap replicates were generated by sampling genes randomly in each category. Genes in each replicate were concatenated and the Grapes program run with the GammaExpo distribution of fitness effect (Galtier 2016). For each recombination category, the mean estimates of α and ω_a_, as well as the standard error over the 100 replicates, were computed.

### Genome-wide analysis of selection patterns

We inferred the strength of purifying selection by computing the pN / pS ratio for each gene. Average pairwise synonymous (πS) and non-synonymous (πN) nucleotide diversity were computed for each genes, and divided by the average number of synonmous (NS) and non-synonymous (NN) positions, respectively, in order to compute the pN / pS ratio as (πN / NN) / (πS / NS). We compared the strength of purifying selection of each gene to several variables, after discarding 6 genes with pN / pS greater than one, as they might be under positive selection. Local recombination rate in 20 kb windows was obtained from Croll et al (Croll *et al.* 2015), and averaged over the two crosses. Population recombination rates (ρ) in the same 20 kb windows were computed as in (Stukenbrock & Dutheil 2017). Each gene was assigned a recombination rate based on the window(s) it overlap with, using a weighted average in case it overlap with multiple windows. Local protein coding site and TE densities were computed as the proportion of coding sites in a window starting x kb upstream and ending x kb downstream each gene. We compared different estimations for x = 10, 20, 50 or 100 kb (see Supplementary Data). For the density of coding sites, we find very little influence of the window size, with a unimodal distribution around ~50%. We therefore selected the intermediate x = 50 kb. The density of TEs showed a large pick at 0 for low values of x. We therefore selected x = 100 kb in order to get a unimodal distribution. GC content at third codon position (GC3) and protein length were also recorded. Expression levels were calculated from (Kellner *et al.* 2014). The mean expression level was computed as the maximum value observed for the gene in axenic culture or plant infection, each averaged over three biological replicates. Genes located on accessory chromosomes were labeled as “dispensable”. Correlation and distribution comparison of the pN / pS ratio with each explanatory variable were performed using rank-based tests (Kendall correlation and Wilcoxon test), as implemented in the R statistical software.

### Estimation of codon usage in Z. tritici

We selected the 10% *Z. tritici* most expressed genes and computed the relative synonymous codon usage of every codons (Sharp *et al.* 1986). Analyses were conducted using the ’uco’ function of the seqinr package for R (Charif *et al.* 2005).

### Model of codon sequence evolution

We used all 9,412 filtered CDS alignments to reconstruct genealogies for the individual genes using PhyML (model HKY85) (Guindon & Gascuel 2003). To investigate patterns of selection and infer the role of positive selection on adaptive gene evolution, the program CodeML from the PAML package was used (Yang 2007) with the filtered multiple CDS alignments and the corresponding phylogenetic trees as inputs. CodeML allows inference of selection and evolutionary rates by calculating the parameter ω, the ratio of non-synonymous to synonymous rates (dN/dS) for each gene. More specifically, we compared site models that allow ω to vary among codons in the protein (Nielsen & Yang 1998). The models used in this study include the nearly neutral (M1a), positive selection (M2a), beta&ω (M8) and bate&ω=1 (M8a) models. A likelihood ratio test (LRT) was used to compare the fit of null models and alternative models, and the significance of the LRT statistic was determined using a χ² distribution. The first LRT tests for the occurrence of sites under positive selection by comparing the M1a and M2a models. In the model M1a sites can be under purifying selection (0 < ω < 1) and evolve by neutral evolution (ω = 0) while the M2a model allows for some sites to be under positive selection (ω > 1). The second LRT compares the M8a and M8 models, where in M8 a discretized beta distribution for ω (limited to the interval [0,1]) and an additional category of sites with ωs > 1. M8a is obtained by constraining ωs > 1 setting (Swanson *et al.* 2003). By allowing for a wider range of strength of purifying selection, the M8 models are more biologically realistic. They may suffer, however, of the same issue than the M7-M8 LRT, which was shown to display an increased false discovery rate compared to the M1a-M2a comparison (Anisimova *et al.* 2001). We corrected for multiple testing and a false discovery rate of 1% was used for the detection of genes under positive selection (Benjamini & Hochberg 1995). Only genes significant for both tests were considered as genes evolving under positive selection (787 out of 9,412 genes analyzed).

To address divergent adaptation, we compared gene evolution among four closely related Zymoseptoria species. In a previous study we defined the core proteome of *Z. tritici*, *Z. ardabiliae*, *Z. brevis* and *Z. pseudotritici* comprising 7,786 orthologous genes (Grandaubert *et al.* 2015). We generated alignments of the corresponding coding sequences using the MACSE sequence aligner (Ranwez *et al.* 2011) and used CodeML with a branch model that allows ω to vary among branches of the phylogeny (Yang & Nielsen 1998). As input we applied a non-rooted tree of the four *Zymoseptoria* species as published in (Stukenbrock *et al.* 2012). Branch lengths were re-estimated for each gene by CodeML.

### Simulation with recombination

We used the coalevol program in order to simulate codon alignments in the presence of recombination (Arenas & Posada 2014). We used a haploid effective population size of 10,000 (option −e10000 1), a one year generation time (option −/1), one parameter for relative transition vs. transversion rate, set to 2 (option −v1 2), a Goldman-Yang model of codon evolution, with 4 omega classes, in equal proportion and set to 0, 0.33, 0.66 and 1.0, respectively (option −m2 4 0.0 0.25 0.33 0.25 0.66 0.25 1). Two mutation rates were tested, 1.10^−5^ and 1.10^−6^ (option -u). One set of simulations was conducted without recombination (-r 0.0), and for others a fixed number of recombination events was used, equal to 1, 2, 5, 10, 30 or 50 (option -w). Protein length was set to 100, 500, 1,000 or 2,000 codon, for 13 and 30 sequence (-s option). Thirty replicates were generated for each parameter combination, and a single phylogenetic tree was inferred using maximum likelihood on the resulting nucleotide aligned, with identical parameters to the real data analysis. M1a and M2a models were then fitted using the estimated tree as input with CodeML. CodeML output was parsed using BioPython (Cock *et al.* 2009).

### Functional enrichment analysis

PFAM domains were extracted from Interproscan results from (Grandaubert *et al.* 2015). Only domain hits with e-values lower than 1.10^−5^ were considered resulting in 10,026 domains present in 7,343 genes. Enrichment tests were performed based on contingency tables, counting the number of genes containing the domain and the number of genes which do not contain it, for both the complete proteome and a given set of candidates to test. A χ² test was performed to assess significance.

### Gene cluster analysis

To analyze the distribution of genes under positive selection, we considered two genes separated by less than 5,000 bp to be clustered and assessed the probability of such clusters under a random distribution of genes along the chromosomes. To do so on a genome-wide scale, we calculated the probability to obtain clusters encompassing from two to ten genes under positive selection when these genes are randomly distributed across all gene coordinates. Based on 10,000 random permutations, it appeared that only clusters containing more than three genes were significant at the 5% level.

### Association between positive selection and effector-encoding genes

To test whether effector-encoding genes are more likely to be under positive selection, we fitted linear models with (1) dN / dS and (2) dN / dS > 1 as response variables, and whether the gene was predicted to encode an effector protein in *Z. tritici* as an explanatory variable. Models were fitted independently for each branch of the four species phylogeny. A binary logistic regression was fitted in order to predict the occurrence of genes under positive selection. For model (1), residues were normalized using a Box-Cox transform as implemented in the MASS package for the R statistical software. An ordinary least square fit was then obtained using the ols function of the rms package for R (Harrell 2015), using the robcov function to obtain robust estimates of the size effects and associated p-values. For model (2), the lrm function of the rms package was used to fit the binary logistic regression model, together with the robcov function to get robust estimates.

### Authors’ contributions

JD and EHS conceived and planned the experiments. JG and JD established the computational framework and analyzed the data. All authors contributed to the interpretation of data and wrote the manuscript. All authors read and approved the final manuscript.

## Acknowledgements

The authors thank Nicolas Galtier and Thomas Bataillon for helpful discussion. The study was funded by a Max Planck fellowship and a personal grant from the State of Schleswig-Holstein, Germany both to EHS. This work was supported by the German Research Foundation (Deutsche Forschungsgemeinschaft), within the priority programs (SPP) 1819 and 1590.

## Supplementary material

**Table S1:** Summary table of isolates used in this study and genome assembly statistics.

**Table S2:** Summary statistics of the multiple genome alignment of thirteen *Z. tritici* genomes.

**Table S3:** Output of the PAML analysis using codon site models for the 9,412 filtered CDS of *Z. tritici*.

**Table S4:** Functional enrichment analysis using PFAM domains for the 787 genes with sites under positive selection in *Z. tritici*.

**Table S5:** Functional enrichment analysis using PFAM domains for the genes under positive selection in four *Zymoseptoria* species.

**Fig. S1:** Codon usage in *Z. tritici*. Relative synonymous codon usage (RSCU) in the 10% most expressed genes of *Z. tritici*. Codon usage, according to the base type at the third position.

